# Long-term temporal stability of circulating proteins in older adults

**DOI:** 10.1101/2025.10.23.684100

**Authors:** Hulda K Ingvarsdottir, Heida Bjarnadottir, Elisabet A Frick, Eva Jacobsen, Kari Arnarson, Nancy Finkel, Joseph J Loureiro, Lenore J Launer, Thor Aspelund, Valur Emilsson, Arnar Palsson, Vilmundur Gudnason, Valborg Gudmundsdottir

**Affiliations:** Faculty of Medicine, University of Iceland, Reykjavik, Iceland; Icelandic Heart Association, Kopavogur, Iceland; Faculty of Pharmaceutical Sciences, University of Iceland, Reykjavik, Iceland; Novartis, Cambridge, MA, USA; Laboratory of Epidemiology and Population Sciences, National Institute on Aging, National Institutes of Health, Baltimore, MD, USA; Institute of Life and Environmental Sciences, University of Iceland, Reykjavik, Iceland

## Abstract

Circulating proteins reflect diverse biological processes and can offer critical insights into an individual‘s overall health and aging trajectory. The circulating proteome is shaped by a complex interplay of genetic, biological, and environmental factors across the lifespan. However, little is known about which factors influence its long-term temporal stability. Here we used SomaScan proteomics to evaluate the five-year temporal stability of 7,288 proteins measured in serum from 3,093 participants (mean age 76 years) of the Age, Gene/Environment Susceptibility (AGES)-Reykjavik study. We observed a wide variability in the temporal stability of individual proteins, with temporally stable proteins more often being extracellular and associated with diseases, while temporally variable proteins are typically involved in intracellular housekeeping functions. We demonstrate that temporal stability of circulating proteins does not reflect that of transcriptomic stability in tissues, and that genetic effects and disease stage are two major contributors to protein temporal stability in the circulation. Our findings underscore the protein-specific differences in long-term temporal stability, and the genetic and biological factors influencing them, which are particularly important to consider in the context of biomarker development and precision medicine.

## Introduction

The circulating proteome, which consists of both actively secreted proteins and those entering the bloodstream following tissue damage^2^, has been intensively studied to understand how protein levels vary between individuals in relation to health and disease, and as a source of novel biomarkers^3^. Recent technological advances have enabled high-throughput measurement of thousands of circulating proteins simultaneously^4^, uncovering links between protein abundance and genetic variation, disease risk, and disease progression. For example, we and others have highlighted the strong influence of genetics on circulating protein levels^5–9^, and proteomic profiling has revealed widespread associations between protein abundance and common diseases^10–17^. We have previously shown that the circulating proteome is organized into co-regulated modules linked to multiple disease states^9^, and a recent study demonstrated that key contributing factors to plasma protein variability include kidney and liver function, smoking, and inflammation^8^. As a collective emphasis has been on using large cohorts to study various disease states, less focus has been put on how protein levels fluctuate within and between individuals over time. Understanding both inter- and intra-individual variability is particularly important in the context of precision medicine, which aims to tailor prevention and treatment strategies to each person’s unique characteristics^18^.

Protein levels have been shown to change with aging^19^, motivating the development of proteomic aging clocks to estimate chronological age and predict mortality risk^20,21^. Longitudinal studies have also revealed nonlinear patterns in proteomic aging, including distinct waves of change in midlife and later life^19,22,23^. Some longitudinal studies have also shown that the circulating proteome as a whole is mainly stable within individuals but displays protein-specific variability. For instance, three studies using different plasma proteomic technologies (Olink^4^, mass spectrometry^24^ (MS), or multiplexed antibody assays^25^), with relatively few proteins and individuals, consistently found that proteomic profiles are highly individual-specific and generally stable, with greater variability between individuals than within the same individuals over time (1-2 years). Stability, however, varied across proteins^4^ and depended on definitions and measurement approaches. Studies integrating proteomic and genomic data further revealed strong genetic contributions to these stable patterns^26^. Collectively, these studies demonstrate the importance of identifying factors that influence dynamics of the circulating proteome that can mirror the real-time global health state of individuals. Despite these insights, prior work has been constrained by small cohorts, short follow-up periods, and limited protein coverage. Furthermore, it is unknown whether temporal stability impacts disease onset in common aging diseases.

To address these limitations, we leveraged data from 3,093 participants from the AGES-Reykjavik cohort^27^, a large-scale, longitudinal, population-based study of elderly Icelandic individuals with roughly 7,000 proteins measured using the SomaScan-v4.1 platform. We aimed to investigate the temporal stability of serum protein levels over two time points, spanning a period of five years, and explore how this stability relates to both technical and biological factors, such as genetic variation and disease states.

## Results

### Temporal stability of serum proteins

The current study included 3,093 AGES participants aged 66–93 years (mean 75 years), of whom 1,794 (58%) were women. Each individual had serum measurements for 7,288 human proteins (SomaScan v4.1) available from two study visits on average 5.2 years apart (Supplementary Table 1), and genotype data available^6^.

Temporal stability of serum proteins between time points was assessed using the Spearman correlation coefficient (hereafter referred to as temporal SCC). Temporal stability varied across proteins, as indicated by a wide distribution of temporal SCC estimates, with a median value of 0.58 (Fig. 1A). Based on suggested guidelines for the interpretation of SCC values^28^, we defined the temporally stable proteins (n_prot_ = 729) as having SCC greater than 0.75 and temporally variable proteins (n_prot_ = 681) less than 0.40, which closely approximated the 90^th^ and 10^th^ percentiles of the distribution (0.75 and 0.41, respectively) (Fig. 1A; Supplementary Table 3). Proteins with intermediate SCC values (0.4–0.75) were classified as having medium temporal stability. This approach ensures that the stable and variable groups represent proteins with clearly distinct temporal behavior, while avoiding arbitrary inclusion of borderline cases.

**Figure 1:**
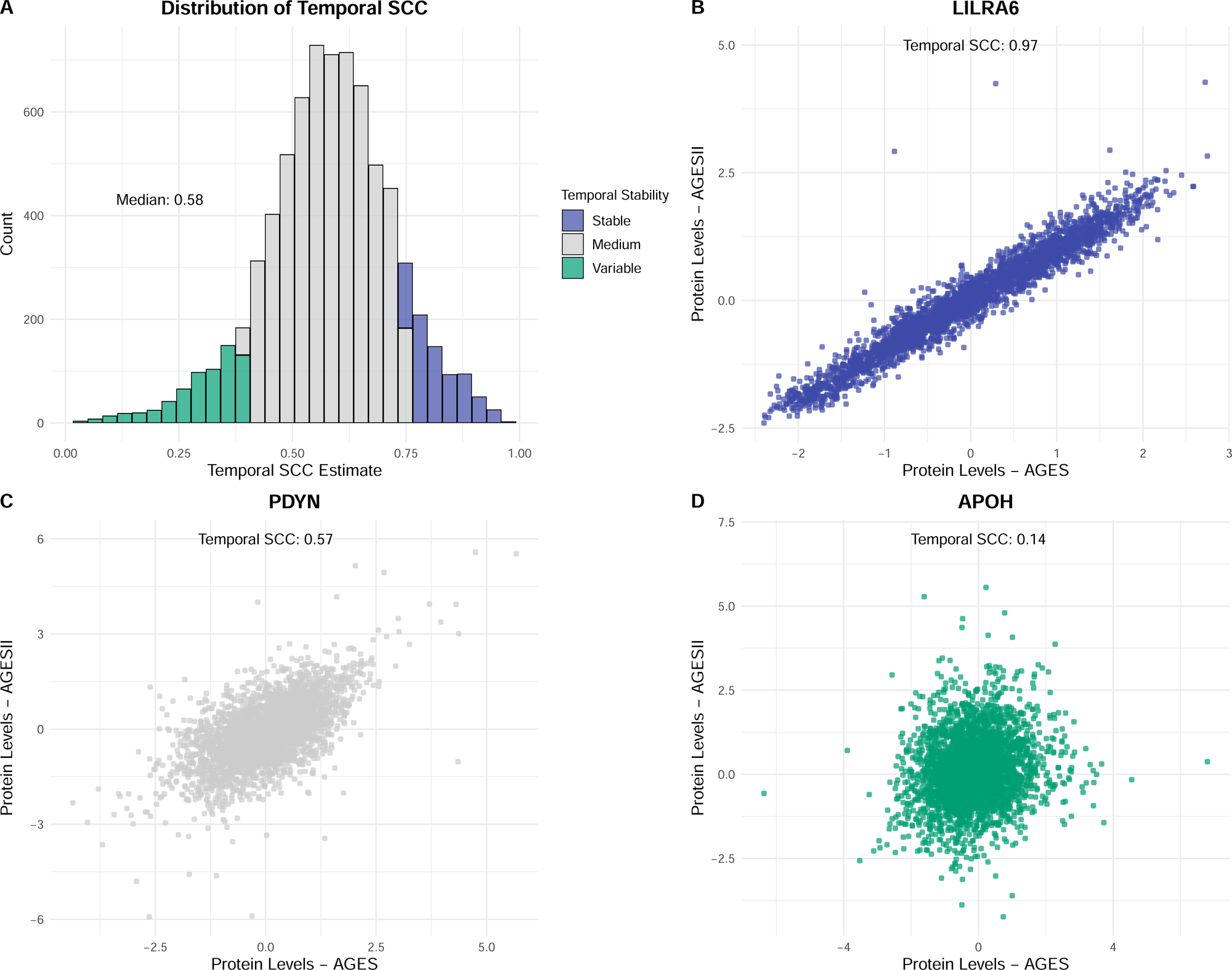
Temporal stability of proteins (n = 7,288). The temporal stability of proteins was assessed using temporal SCC estimates. Proteins were divided into three groups: temporally variable (<0.40), medium (0.40–0.75), and stable (>0.75). **A)** Distribution of temporal SCC estimates of all proteins. **B – D)** Examples of proteins from each group, showing baseline (AGES) versus follow-up (AGESII) protein levels for each individual. **B)** LILRA6 was the most temporally stable protein (temporal SCC = 0.97). **C)** PDYN showed medium temporal stability (temporal SCC = 0.57). **D)** APOH was temporally variable (temporal SCC = 0.14).

The most temporally stable protein was LILRA6 (temporal SCC = 0.97; Fig. 1B), a leukocyte immunoglobulin-like receptor A6 that functions as a receptor for MHC I antigens^29^. An example of a protein with medium temporal stability was PDYN (prodynorphin; temporal SCC = 0.57; Fig. 1D), a precursor of opioid peptides that modulate pain, stress, and mood through the κ-opioid receptor system^30^. Finally, an example of a highly temporally variable protein was APOH (apolipoprotein H; temporal SCC = 0.14; Fig. 1D), a glycoprotein that binds to phospholipids and other negatively charged molecules in the circulation and has been linked to systemic lupus erythematosus (SLE) and antiphospholipid syndrome^31^.

The temporal SCC showed almost no correlation with the absolute difference in protein measurements (median delta value) between time points (SCC = 0.05; 95% CI: 0.03-0.08; p-value = 3.90 × 10^−6^; Supplementary Fig. 1). Although statistically significant, the effect size was negligible, indicating that the correlation is not practically meaningful. This suggests that temporal SCC captures a complementary aspect of protein temporal stability rather than simply reflecting absolute changes in protein levels over time. However, there was a moderate relationship between the intra-individual variability (temporal SCC) and inter-individual variability of proteins, both at baseline (AGES) and the follow-up (AGESII) visit (SCC AGES = 0.22, SCC AGESII = 0.26), where proteins with higher intra-individual variability were also more likely to have high inter-individual variability (Supplementary Fig. 2).

We tested whether temporal stability was associated with relative protein abundance in AGES, as measured by their median relative fluorescent unit (RFU) values, but relative protein abundance showed a strong correlation with external blood concentration estimates from the Human Protein Atlas (SCC = 0.71, p < 2.2 × 10^−16^; Supplementary Fig. 3), available for 569 proteins. Pairwise comparisons between temporal stability protein groups revealed that the stable proteins had significantly higher protein abundance than those in the medium or variable groups (Supplementary Fig. 4).

To estimate whether our results reflect technical variability, we extracted inter-plate percentile variation (PV) values from a recent reproducibility study of the SomaScan v4.1 platform conducted in the Atherosclerosis Risk in Communities (ARIC) Study^32^. We compared these PV values with the temporal SCC value in AGES for each protein. As expected, this comparison revealed a negative relationship where proteins with lower temporal SCC estimates in AGES tended to have higher PV in ARIC, indicating more technical variability among the proteins defined as temporally variable (Supplementary Fig. 5). Roughly 82% of the temporally variable proteins had a PV < 14.11% in ARIC, corresponding to Z-scores ≤ 2, indicating that they are technically robust. Proteins with PV > 14.11% (Z-score > 2) were excluded from downstream analyses to reduce the effect of technical variability when comparing stable and variable proteins, leaving 7,097 proteins for analysis. We note that while abundance generally correlates with temporal stability, exceptions exist, including low-abundance proteins that are temporally stable and high-abundance proteins that exhibit temporal variability.

### Functional characteristics of temporally stable and variable proteins

To evaluate the characteristics of the temporal stability protein groups, we performed functional enrichment analysis using all SomaScan 7k proteins as the background. Overall, a clear functional separation was observed between the stable and variable protein groups when incorporating GO terms, KEGG, WikiPathways, and Reactome (Fig. 2A; Supplementary Table 4), indicating a biological distinction relating to temporal stability. Only a few terms were significantly associated with both stable and variable proteins, including the immune system and the secretory granule. The terms “extracellular region” and “extracellular space” were significantly enriched among both temporal stability groups, although this enrichment was much stronger among temporally stable proteins than among the variable group.

**Figure 2:**
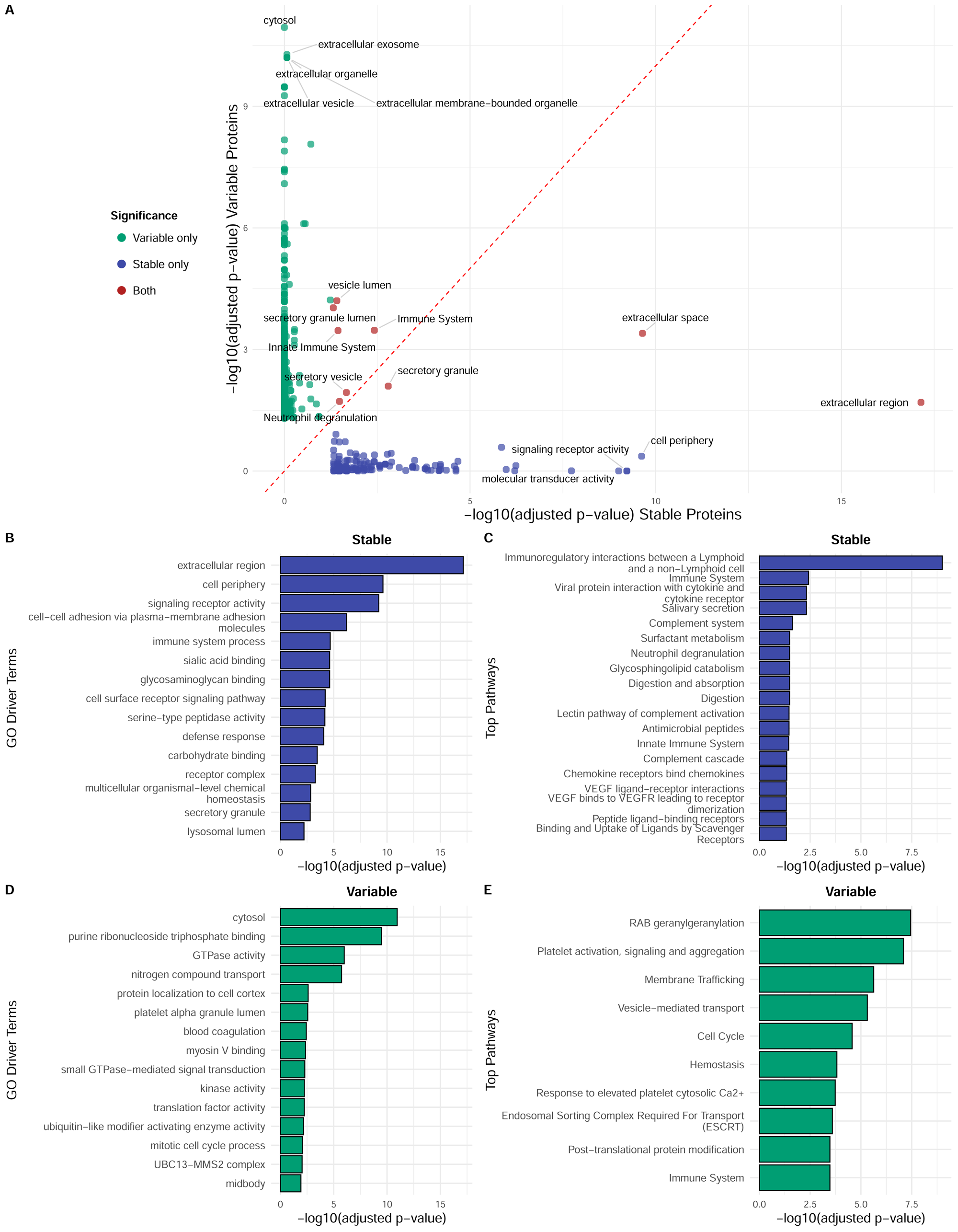
Enrichment and functional analysis using g:Profiler. **A)** Comparison of enriched pathways between temporally stable and variable protein groups, showing distinct biological roles with a few pathways shared across groups. Data points are colored based on whether temporally stable, variable proteins, or both significantly enriched the pathway. **B)** Top 15 enriched Gene Ontology (GO) driver terms for temporally stable proteins. **C)** Enriched pathways for temporally stable proteins from KEGG, Reactome, and WikiPathways. **D)** Top 15 GO driver terms for temporally variable proteins. **E)** Enriched pathways for temporally variable proteins from KEGG, Reactome, and WikiPathways.

For the temporally stable proteins, the gene ontology (GO) driver terms with the strongest enrichment included extracellular region (FDR = 7.18 × 10^−18^), cell periphery (FDR = 2.40 × 10^−10^), and signaling receptor activity (FDR = 5.99 × 10^−10^, Fig. 2B). The strongest enriched pathways were observed in immunoregulatory interactions between a lymphoid and a non-lymphoid cell (FDR = 9.92 × 10^−10^), immune System (FDR = 3.78 × 10^−3^), viral protein interaction with cytokine and cytokine receptor (FDR = 4.85 × 10^−3^), salivary secretion (FDR = 4.85 × 10^−3^), complement system (FDR = 2.31 × 10^−2^), and digestion and absorption (FDR = 3.28 × 10^−2^, Fig. 2C). Other highly enriched pathways include glycosphingolipid catabolism, digestion, neutrophil degranulation, and surfactant metabolism (Supplementary Table 5). Thus, temporally stable proteins tend to have established extracellular roles, particularly in signaling and immune pathways.

In contrast, the variable proteins were most strongly enriched for the GO driver terms cytosol (FDR = 1.13 × 10^−11^), purine ribonucleoside triphosphate binding (FDR = 3.28 × 10^−10^), and GTPase activity (FDR = 1.02 × 10^−6^, Fig. 2D). The top pathways enriched in the variable protein group included RAB geranylgeranylation (FDR = 3.55 × 10^−8^), platelet activation, signaling and aggregation (FDR = 8.10 × 10^−8^), membrane trafficking (FDR = 2.34 × 10^−6^), vesicle-mediated transport (FDR = 4.85 × 10^−6^), cell cycle (FDR = 2.71 × 10^−5^), and hemostasis (FDR = 1.53 × 10^−4^, Fig. 2E). Additional enriched pathways included response to elevated platelet cytosolic Ca2+, Endosomal Sorting Complex Required For Transport (ESCRT), immune system, and post-translational protein modification (Supplementary Table 6). Overall, the temporally variable proteins are thus more likely to participate in intracellular functions related to cell cycle regulation and intracellular trafficking.

Mirroring the patterns observed from GO term and pathway memberships, the temporal SCC protein groups were also enriched for different protein classes, as defined by the Human Protein Atlas (Fig. 3A). The stable proteins were significantly enriched for predicted secreted proteins, predicted membrane proteins, CD markers, and plasma proteins. In contrast, the variable proteins were enriched for predicted intracellular proteins, enzymes, cancer-related genes, and plasma proteins. Furthermore, stable proteins were more likely to be encoded by tissue-enriched genes and to exhibit tissue-enriched protein expression, whereas temporally variable proteins were under-represented in both categories. Finally, the temporally variable proteins were less likely to tolerate loss-of-function (LoF) mutations (Fig. 3B) and had significantly higher hub status in both a physical protein-protein interaction (PPI) network^33^ (Fig. 3C; Wilcoxon test, two-sided, p < 1 × 10^−4^) and the serum co-regulatory network^9^ (Fig. 3D; Wilcoxon test, two-sided, p < 1 × 10^−4^).

**Figure 3:**
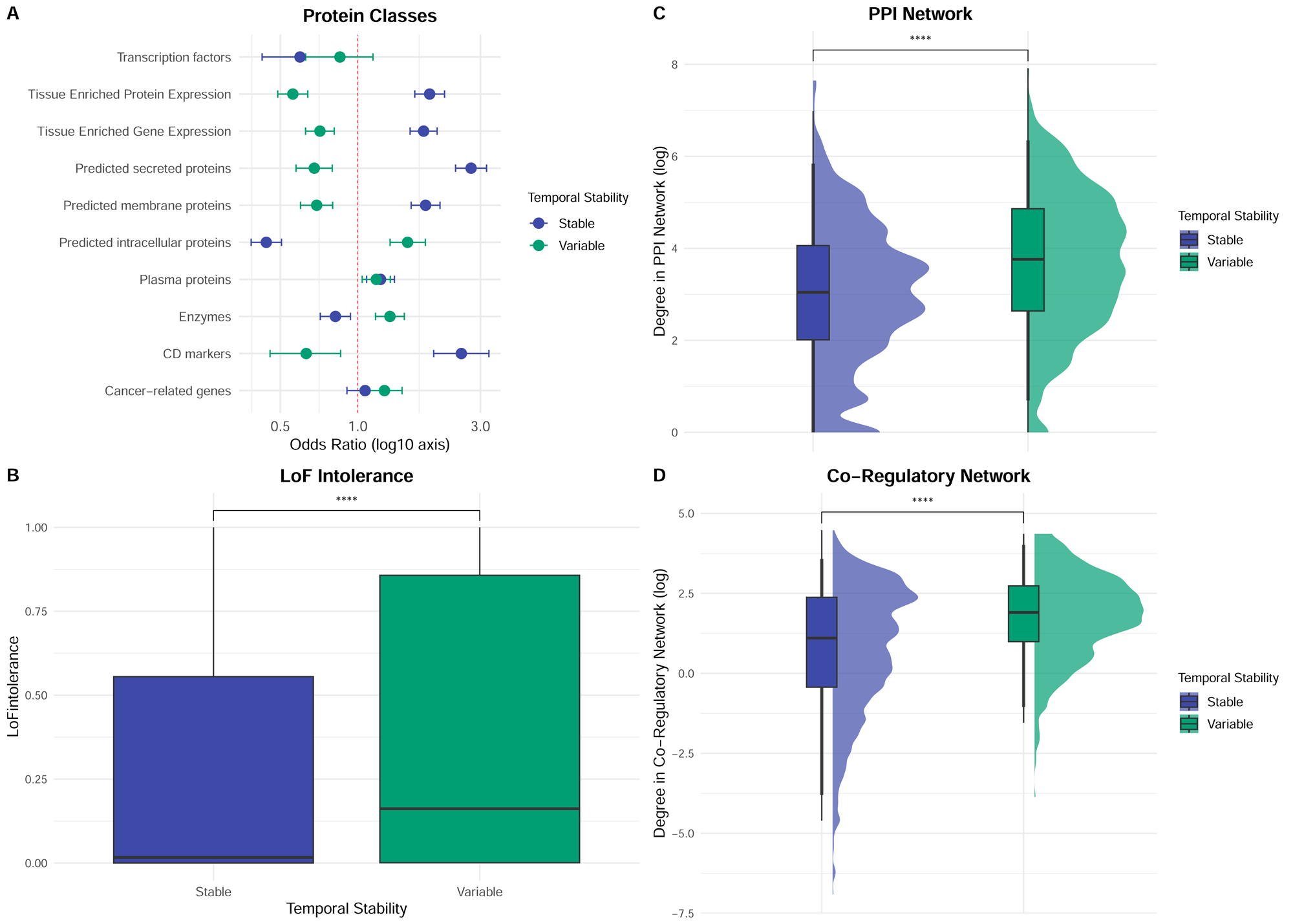
Functional characteristics of temporally stable and variable proteins. **A)** Protein classes and tissue-enriched gene/protein expression (x-axis log-scaled). Odds ratio estimates are presented with 95% confidence intervals. **B)** Loss-of-function (LoF) intolerance, with variable proteins showing significantly higher LoF intolerance (Wilcoxon test, two-sided, p < 1 × 10□³). **C)** Degree (log-scaled) in the protein–protein interaction (PPI) network, where variable proteins had significantly higher connectivity, indicating they are more likely to be hubs. **D)** Degree (log-scaled) in the co-regulatory serum network, where variable proteins also showed higher connectivity (Wilcoxon test, two-sided, p < 1 × 10□³). Boxplots in **(B–C)** indicate median value, 25^th^ and 75^th^ percentiles. Whiskers extend to smallest/largest value of no further than 1.5 X interquartile range. Outliers are not shown.

### Gene expression variance and protein temporal stability

The enrichment results above demonstrated that the temporally variable proteins were generally more involved in housekeeping processes than the stable protein group. This was somewhat unexpected, as our initial hypothesis was that housekeeping proteins would have stable expression and, thus, be temporally stable in the longitudinal serum proteomics data. We therefore explored gene expression variability over time and its relationship with circulating protein temporal stability to evaluate if this pattern would also be reflected at the transcriptomic level and across different tissues.

We utilized data from a study examining the landscape of gene expression variance across 57 RNA-sequencing datasets from various tissues and distinct populations^34^. In this study, gene expression variance was calculated as the standard deviation (SD) of gene expression levels for each gene across all samples, with each gene ranked based on this value. We observed a significant difference in gene expression variance between the temporally stable and variable protein groups (Wilcoxon test, two-sided, p < 1 × 10^−3^; Supplementary Fig. 6). Specifically, the stable proteins had higher SD rank values, indicating more gene expression variance overall, whereas variable proteins had lower gene expression variance. This suggests an inverse relationship between gene expression variance and protein temporal stability. Overall, this observation indicates that temporal stability in serum may only partially capture the transcriptomic variance across tissues and individuals and may instead reflect the mechanisms by which the proteins enter, or are cleared from, the circulation.

### Genetic effects on temporal stability

We have previously described the genetic effects on proteomic measurements in the AGES study^6,35^. In total, 6,572 SOMAmers (targeting 5,798 unique proteins) had at least one conditionally independent and genome-wide significant (p < 5 × 10^−8^) protein quantitative trait locus (pQTL). Among these, 278 SOMAmers had only cis-pQTLs (residing within 500 kb of the respective gene), 1,468 had both cis- and trans-pQTLs (hence not direct transcript variants), and 4,826 had only trans-pQTLs. The remaining 716 SOMAmers had no significant pQTLs.

To assess whether genetic signals influence protein expression over time, all 7,288 protein measurements (see Methods) were adjusted for conditionally independent pQTLs, therefore removing genotype variability in the protein measurements. The temporal SCCs were then recalculated (hereafter referred to as *adjusted temporal SCC*) to assess any shifts in temporal stability upon the removal of genetic drivers. The comparison between original and adjusted temporal SCC values (Fig. 4A) shows a strong overall correlation (SCC = 0.94), and the median SCC changed from 0.58 to 0.56 (paired t-test, two-sided, p ≤ 1 × 10^−3^).

**Figure 4:**
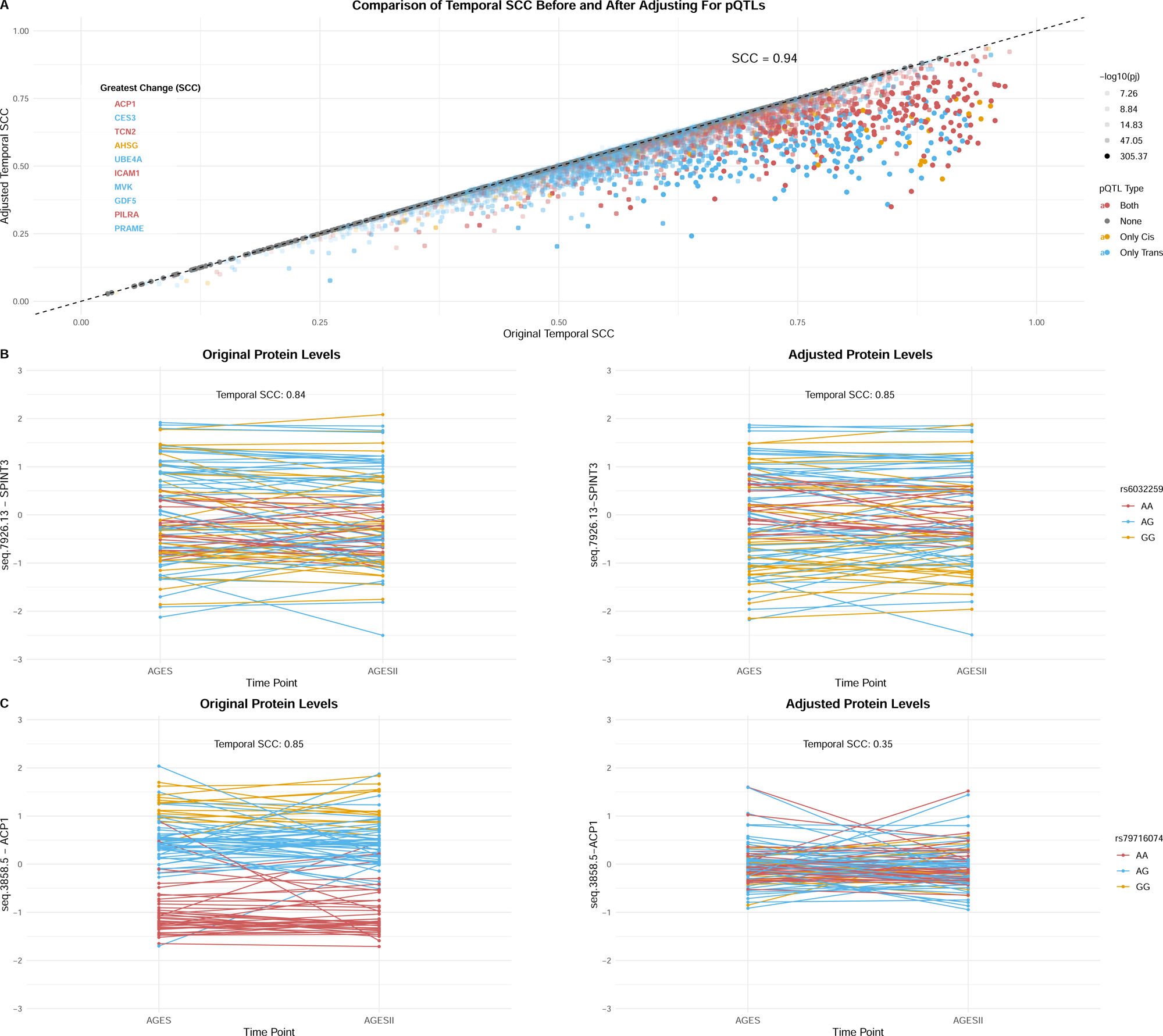
Effect of strong pQTLs on temporal stability. **A)** Scatter plot of temporal SCC estimates before and after adjusting protein levels for conditionally independent pQTLs (p < 5 × 10□□). Original and adjusted SCC values were strongly correlated (SCC = 0.94), but 75% of SOMAmers showed lower SCC values after adjustment. Proteins are colored by pQTL type, with less significant points shown at reduced opacity. **B–C)** Examples of proteins initially classified as temporally stable. **B)** SPINT3 remained temporally stable after adjustment (original SCC = 0.84; adjusted SCC = 0.85). **C)** ACP1 (original SCC = 0.85) initially showed genotype-based separation before adjustment but was reclassified as temporally variable after adjustment (SCC = 0.35), with reduced genotype separation and more apparent within-individual fluctuation.

Nevertheless, following adjustment, 75% of SOMAmers exhibited lower SCC values, leading to a subset of 5 proteins previously classified as stable being redefined as variable over time. The ten proteins with the largest decrease in temporal SCC (ranging from 0.39 to 0.50) were ACP1, CES3, TCN2, AHSG, UBE4A, ICAM1, MVK, GDF15, PILRA, and PRAME. This decrease was primarily driven by only trans- or both cis and trans signals, where all except one protein, AHSG (only cis), had more than one independent pQTL signal (Fig. 4A). In other words, these proteins had more variable protein expression over time when genetic effects had been accounted for.

Many of these proteins are influenced by strong genetic signals and have been implicated in various biological processes and disease mechanisms. For instance, GDF15 has been associated with multiple conditions, including cancer, cardiovascular disease, and obesity,^36^ and we have previously shown suggestive evidence that higher circulating GDF15 levels may causally increase the risk of type 2 diabetes^11^. PILRA has also been proposed as a potential therapeutic target for Alzheimer‘s disease^37^ and atrial fibrillation^38^. Overall, these results underscore the need to consider genetic contributions when evaluating protein temporal stability, which may provide novel insights into disease mechanisms and potential therapeutic targets.

SOMAmers with cis-pQTLs had higher temporal SCC both before and after adjustment (Supplementary Fig. 7A). There was also a significant difference between original and adjusted temporal SCC across all pQTL groups except for proteins with no pQTL signal, and proteins with both cis- and trans-pQTLs showed the highest difference (Supplementary Fig. 7B). These findings suggest that proteins under strong cis-acting genetic regulation tend to be more temporally stable. In contrast, proteins lacking such pQTL signals may be more influenced by external or environmental factors, leading to greater variability over time. This aligns with our previous results, which state that temporally variable proteins are more likely to be hubs and protected against functionally annotated variants (see Functional characteristics of temporally stable and variable proteins).

Notably, some proteins remained temporally stable despite the adjustment for strong pQTLS. For example, SPINT3 (serine peptidase inhibitor, Kunitz type 3), associated with a strong cis-pQTL (rs6032259, conditional p = 1.01 × 10^−93^), maintained stability (temporal SCC = 0.84 before vs. 0.85 after adjustment; Fig. 4B). By contrast, ACP1 (acid phosphatase 1), originally stable (temporal SCC = 0.85) and associated with a strong cis-pQTL (rs79716074, conditional p < 1 × 10□³□□), showed a pronounced decrease in stability after adjustment (temporal SCC = 0.35). Moreover, ACP1 exhibited high variability in protein levels within each genotype and between time points, suggesting that its apparent stability was largely genotype-driven. Together, these results indicate that strong genetic effects on protein abundance can create the appearance of temporal stability, which again underscores the importance of accounting for genetic drivers when interpreting longitudinal protein profiles.

### Temporal stability of co-regulatory protein modules

To evaluate the temporal stability of proteins at the level of co-regulated protein modules, we applied weighted gene co-expression network analysis (WGCNA) to define 25 co-expressed protein modules at baseline and 23 modules at the follow-up visit in the 3,093 individuals with longitudinal measurements (Supplementary Tables 7–8). The co-regulatory modules were enriched for different biological functions (Supplementary Table 9). We compared the baseline modules to the previously constructed co-regulatory network from the AGES cohort using the 5k version of the SomaScan assay in the full AGES baseline sample (n = 5,457)^9^. The overall structure was similar, although some distinct clustering patterns were observed in this subset of individuals (Supplementary Fig. 8).

Integrating information on protein temporal stability, we identified 11 baseline modules enriched for stable proteins, 10 for variable proteins, and 4 for neither (Wilcoxon rank-sum test, two-sided, p < 0.05; Supplementary Table 9, referred to as TS enrichment). An example of a module enriched for temporally stable proteins is M16 (n_prot_ = 192), which is composed of proteins localized to mitochondrial outer membranes, while M8 (n_prot_ = 300) is an example of a module enriched for temporally variable proteins, involved in coagulation cascades (Fig. 5A; Supplementary Table 9).

**Figure 5:**
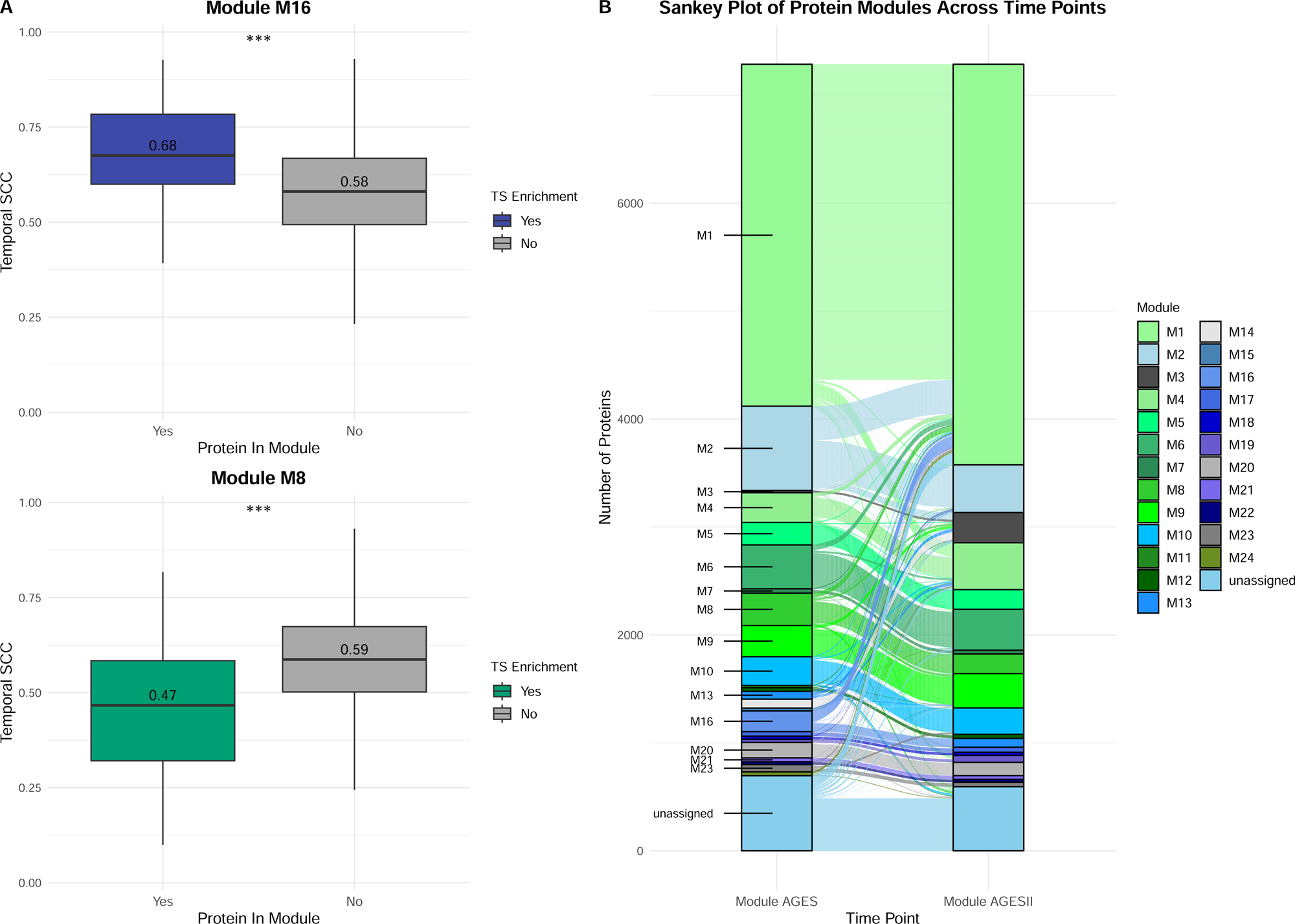
Baseline protein module enrichment for temporal stability protein groups. **A)** Examples of modules enriched for each temporal stability (TS) group. Proteins in module M16 had significantly higher median temporal SCC estimates compared to other proteins (Wilcoxon test, two-sided, p < 1 × 10□³). In contrast, proteins in module M8 had lower median temporal SCC estimates compared to other proteins (Wilcoxon test, two-sided, p < 1 × 10□³). Boxplots indicate median value, 25^th^ and 75^th^ percentiles. Whiskers extend to smallest/largest value of no further than 1.5 X interquartile range. Outliers are not shown. **B)** Sankey plot showing how proteins are grouped into modules at each time point. Some proteins remain in the same modules over time, while others shift between modules. Modules enriched for variable proteins are colored green, stable-enriched modules are colored blue, and modules without significant enrichment are colored grey.

We next explored the longitudinal behavior of co-regulatory patterns by examining the flow of proteins between baseline and follow-up modules (Fig. 5B). Overall, 5,738 (79%) of the 7,288 proteins remained within their dominant module group at follow-up, demonstrating a consistent co-regulatory pattern over time. While most modules retained a consistent protein composition over time, a few exhibited notable reorganization, with proteins shifting to different modules at follow-up. For example, 108 of the 192 proteins (∼56%) in the stable-enriched module M16 moved to module M1 at the follow-up visit (Supplementary Table 10). In comparison, 173 of the 300 proteins (∼58%) in the variable-enriched module M8 were retained in M8 at the follow-up visit. There was no statistically significant difference in protein retention by temporal stability enrichment (stable-enriched modules, 78.6%; variable-enriched modules, 73.9%; no enrichment, 83.1%; *Kruskal-Wallis* _χ_*²* = 0.61; *df* = 2, *p* = 0.74). Thus, our results indicate that temporal stability in circulating protein abundance does not translate into the stability of inter-protein relationships.

Taken together, these observations suggest that while the majority of protein co-regulatory modules are preserved over time and across assay platforms, a subset of modules exhibits temporal or context-dependent variability, highlighting dynamic aspects of the plasma proteome.

### Protein temporal stability in the context of aging and disease

We examined whether proteins classified as temporally stable or variable were more likely to be associated with a range of disease phenotypes and overall mortality in the AGES cohort. We observed significant (FDR < 0.05) associations between baseline protein levels and all tested phenotypes, across both stable and variable proteins. For most diseases, a greater proportion of the associated proteins were temporally stable than variable (Fig. 6A; FDR < 0.05). However, variable proteins were also linked to all phenotypes examined, with three related prevalent diseases (AF, stroke, and all-cause dementia) being associated with a significantly higher proportion of variable proteins (Fig. 6A).

**Figure 6:**
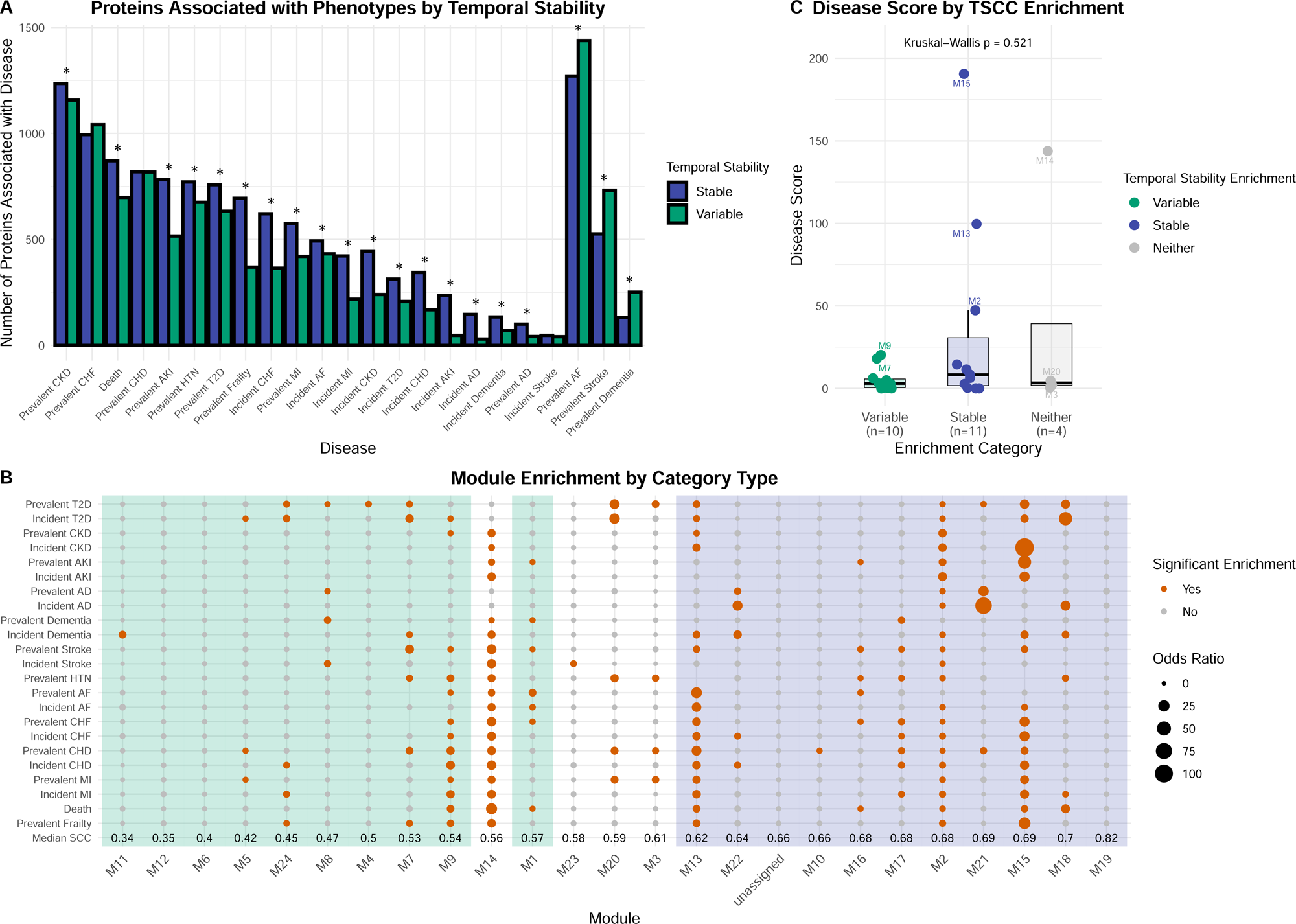
Temporal stability and phenotype associations. **A)** Number of proteins measured at baseline that are significantly associated with each phenotype (excluding proteins with “medium” temporal stability), shown for both incident and prevalent disease outcomes. Bars are colored by protein temporal stability. An asterisk (*) above a phenotype indicates significant enrichment (FDR < 0.05) of either temporally stable or variable proteins associated with that phenotype. **B)** Modules labeled on the x-axis, ordered by median temporal SCC. The y-axis lists each phenotype tested for associations with baseline protein levels. Orange dots indicate significant enrichment for a phenotype within a module, with dot size reflecting the strength of enrichment (odds ratio). Modules are highlighted vertically, with green indicating temporally variable-enriched and blue indicating temporally stable-enriched. **C)** Disease score by temporal SCC enrichment of modules. The score was calculated as the number of associated phenotypes multiplied by the median odds ratio. No significant difference was observed between enrichment categories (Kruskal–Wallis p = 0.54), though some stable-enriched modules exhibited high scores. Boxplots indicate the median, 25th, and 75th percentiles; whiskers extend to the smallest and largest values within 1.5× the interquartile range. Outliers are not shown. All data points are displayed with jitter, and labeled points indicate top modules.

Although temporally stable proteins were generally more likely to associate with disease phenotypes than variable proteins, many proteins are correlated or functionally linked, meaning that interpreting associations individually can be misleading^39^. Taking into account the co-regulatory structure of the circulating proteome and building on our previous analysis of protein modules, we examined whether diseases were enriched within these co-regulatory modules at baseline. Both stable- and variable-enriched baseline modules were enriched for disease associations (Fig. 6B) and comparing disease scores across modules stratified by temporal stability enrichment (see Methods) revealed no significant difference (*Kruskal-Wallis* _χ_*²* = 1.31, *df* = 2, *p* = 0.52; Fig. 6C). Despite this, we identified several stable modules with particularly high disease scores, including M15 (n_prot_ = 26, fatty acid and lipid metabolism), M13 (n_prot_ = 72, RNA metabolism), and M2 (n_prot_ = 781, cell adhesion and extracellular interactions) (Fig. 6B–C). Further information about the characteristics of each baseline module is available in Supplementary Table 9.

To explore the potential effect of disease onset on protein temporal stability, we examined baseline protein modules to determine whether individuals who developed a given disease between visits exhibited different temporal SCC patterns compared to those who did not. We identified three modules containing disease-associated proteins with a significant (empirical P < 0.01) difference in median temporal SCC in individuals diagnosed between the AGES baseline visit (A1) and follow-up (A2), compared to each of the other three groups (diagnosed before A1, after A2, or never diagnosed) (Fig.7). This pattern suggests potential shifts in protein temporal stability in individuals before or during disease onset.

**Figure 7:**
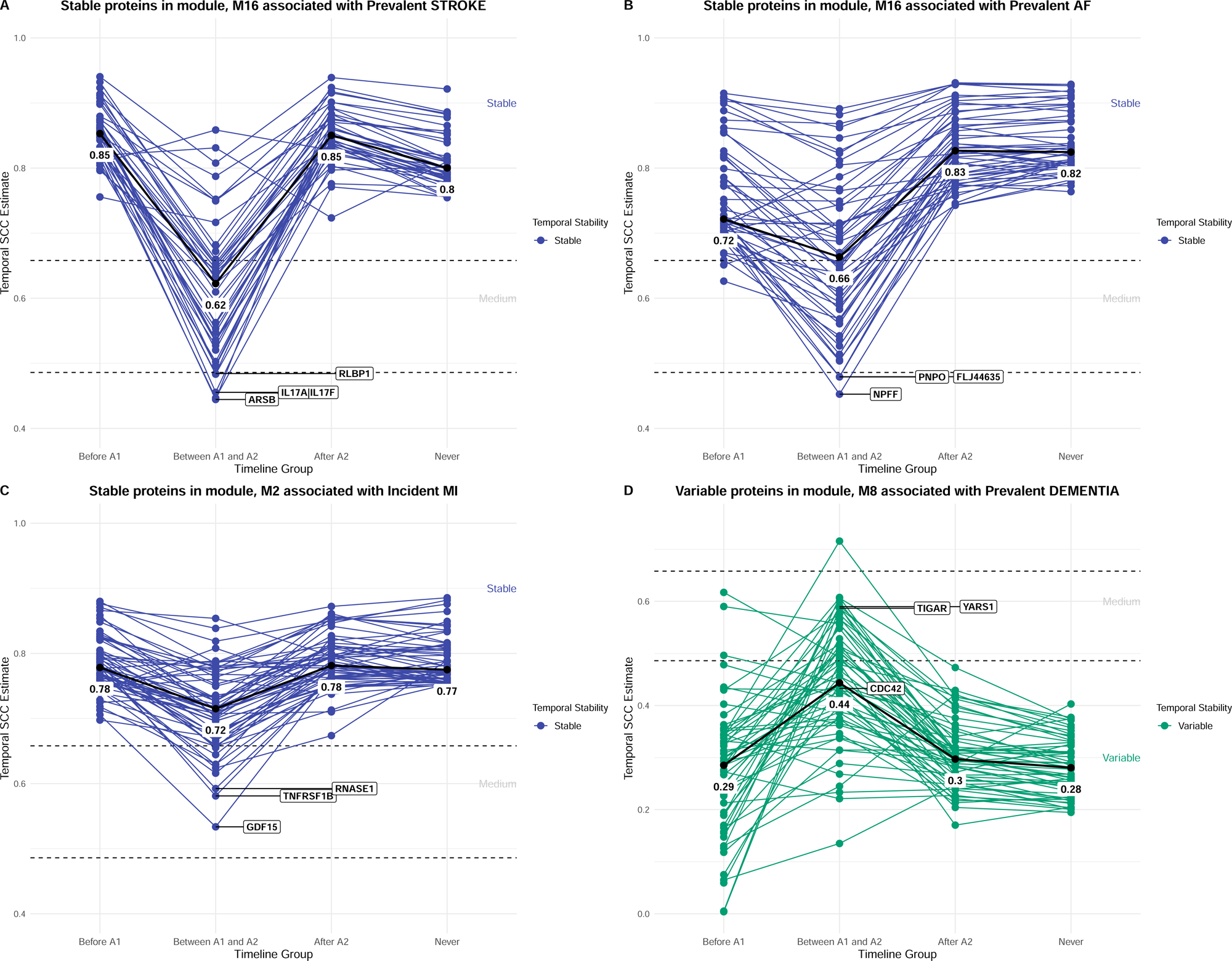
Effect of temporal SCC on disease progression. **A–D)** Three modules included protein groups associated with disease that differed in temporal stability depending on disease stage. For individuals diagnosed between visits (between A1, the baseline visit, and A2, the follow-up visit), there was a significant difference in median temporal SCC (empirical p < 0.01) compared to the other groups: those diagnosed before A1, those diagnosed after A2, or those who were never diagnosed. Two modules were temporally stable-enriched: M16 **(A–B)** and M2 **(C)**, both of which showed reduced temporal SCC in the between-visits group. The temporally variable-enriched module M8 **(D)** showed the opposite pattern, with increased temporal SCC in individuals diagnosed between visits.

Within module M16, stable proteins associated with prevalent AF or stroke showed much lower temporal SCC in individuals diagnosed between A1 and A2 (median = 0.66 and 0.62, respectively) compared to the other timeline groups (median ≥ 0.72 and 0.8, respectively) (Fig. 7A–B). The proteins contributing most strongly to these differences were ARSB, IL17A|IL17F, and RLBP1 for stroke, and NPFF, PNPO, and FLJ44635 for AF. A similar pattern of reduced temporal SCC in individuals diagnosed with incident MI between time points was observed in module M2 (Fig. 7C). The proteins with the largest differences in this module were GDF15, TNFRSF1B, and RNASE1. Finally, variable proteins within module M8 associated with prevalent dementia showed a reversed pattern, as individuals diagnosed between visits had higher temporal SCC (median = 0.44) compared to all other groups (median ≤ 0.3) (Fig. 7D). The proteins most strongly driving this reversed pattern were YARS1, TIGAR, and CDC42. Thus, these findings illustrate how the onset of a disease state can influence the temporal stability of proteins in either direction.

## Discussion

In this study, we investigated the longitudinal variability of serum proteins over a five-year period in older individuals to characterize their temporal dynamics and identify the factors contributing to their temporal stability. Our findings reveal a wide variability in the temporal stability of individual proteins and demonstrate clear biological differences between proteins that tend to remain stable in circulation over time versus those that vary, suggesting different mechanisms of entry into the bloodstream. We also found that both genetic factors and disease onset can influence protein temporal stability, which is particularly important to consider in disease biomarker research.

We identified key biological differences in the characteristics of the temporal stability protein groups, as stable proteins were more likely to be extracellular, secreted, or transmembrane, exhibit tissue-enriched gene and protein expression, be more abundant, tolerate LoF mutations, and have a cis-acting pQTL. In contrast, variable proteins were more likely to be intracellular, enzymes, less tissue-specific, interact with many proteins, and be intolerant to LoF mutations.

Although variable proteins tended to be more intracellular, they were also enriched for pathways such as extracellular exosome, extracellular organelle, extracellular membrane-bounded organelle, and extracellular vesicle. Thus, although these proteins are annotated as part of extracellular pathways or components, they may primarily reflect intracellular proteins that can be found within extracellular vesicles secreted outside the cell^40^. Somewhat surprisingly, we observed that temporally variable proteins in the circulation were more likely to have biological properties consistent with housekeeping genes, which are known to exhibit low variability in expression across tissues and conditions^41^. Our comparison with external transcriptomic data revealed that variable proteins were more likely to have low gene expression variance, whereas stable proteins showed the opposite. This pattern suggests that protein temporal stability in circulation is mainly driven by post-transcriptional regulation, release dynamics, or clearance, rather than reflecting transcriptomic regulation. Overall, our findings provide important insights into the pathways and functions of circulating proteins that differ in relation to their behavior over time.

A previous study examining the longitudinal variability of proteins over one year in 101 clinically healthy individuals demonstrated that each individual had a unique protein profile, largely explained by genetic associations^25^. Another study linked 90-95% of 1.3K SomaScan proteomic profiles to their corresponding genome, highlighting the strong genetic influence on protein levels resulting in an individual proteomic fingerprint^42^. Consistent with these findings, but examining individual protein patterns, our study shows that genetic variability significantly influences the temporal stability of serum proteins. Adjusting for pQTLs decreased temporal stability for 75% of proteins and, notably, some proteins initially classified as stable were even reclassified as variable, suggesting that genetic factors contribute substantially to their stability over time. Such genetic effects are particularly important to consider for biomarker candidates, as they both mask population-level disease associations and person-specific deviations from their baseline over time.

At the levels of co-regulatory protein modules, we found that of all 7,288 proteins, approximately 80% remained in the same module over the five-year study period, while ∼20% shifted to different modules, suggesting that while the co-regulatory structure of the circulating proteome is largely stable over time, there is also dynamic reorganization of protein co-regulatory patterns. This balance between stability and change in co-regulatory networks may reflect underlying biological processes related to aging. As aging is associated with a gradual decline in proteostasis and accumulation of molecular damage affecting protein function and interactions^43^, some proteins likely maintain stable co-regulation due to preserved cellular homeostasis, whereas others shift modules because of age-related dysregulation and proteomic remodeling. Longitudinal module retention was not related to the temporal stability of individual proteins, demonstrating that temporally variable proteins can co-vary and maintain stable inter-protein relationships.

Circulating proteins generally need low biological variability within and between individuals to be useful as clinical biomarkers^44^. Although we did not find a consistent relationship between protein intra- and inter-individual variability, temporally stable proteins were generally more likely to be associated with disease outcomes than variable proteins. While this general pattern was not reflected at the protein module level, the strongest disease enrichment was still observed for the stable modules. This information, together with the stable proteins’ extracellular nature and tissue-specific expression, makes them valuable candidates for drug targets and disease biomarkers^45^, leading us further to interrogate the effects of incident disease manifestation on temporal stability. We proposed that proteins generally stable over time in individuals may exhibit temporally variable patterns in the context of incident disease manifestation and found such examples for proteins associated with stroke, AF, and MI. Specifically, individuals diagnosed between visits showed markedly greater temporal variability in protein levels compared to others. Interestingly, we also identified a group of temporally variable proteins associated with prevalent dementia for which the pattern was reversed, with the protein levels being more stable in individuals between visits. Thus, the temporal stability of specific proteins can be either increased or decreased during the process of disease development, and further studies will be required to elucidate the underlying mechanisms for specific protein candidates.

The proteins in module M16 showing decreased stability at the onset of AF and stroke were enriched for mitochondrial outer membrane proteins (Supplementary Table 9), a cellular component known to regulate apoptotic signaling pathways^46^. Mitochondrial dysfunction is also a well-established contributor to the development of cardiovascular diseases^47^. One of the proteins showing the greatest change in temporal SCC in individuals who developed stroke between time points was IL17A|IL17F, which has previously been linked to stroke-related neuroinflammation^48^. Module M2 comprised proteins linked to incident MI whose stability similarly declined in individuals with events, suggesting that their temporal variability may mark early or subclinical processes common to diverse disease trajectories. This module was enriched for proteins involved in cell adhesion and extracellular interactions (Supplementary Table 9), suggesting that this pattern may indicate long-term dysregulation in these processes, with temporal variability between time points reflecting disease progression.

Finally, module M8 included temporally variable proteins whose baseline values were associated with prevalent dementia but exhibited greater stability in individuals diagnosed between visits. Analysis of this module revealed enrichment for coagulation proteins (Supplementary Table 9), which have been shown to differ in plasma between individuals with AD and healthy controls^49^. One of the top proteins, exhibiting the greatest differences in temporal SCC in individuals diagnosed between visits compared to all other groups, was TIGAR. This protein has multiple physiological functions related to mitochondrial functions, apoptosis, autophagy, anti-oxidative stress, and inflammation^29^. It plays an important role in nervous system diseases and has been proposed as a potential therapeutic target^50^.

Together, these findings suggest that proteins in M8, many of which are involved in coagulation, are typically temporally variable in the general population, even in individuals with prevalent dementia. However, during a particular window of time at the onset of disease, these proteins appear to become more stable. This could reflect a shift toward a new equilibrium in protein expression once the disease process has advanced to a clinically detectable stage, potentially due to compensatory mechanisms^51^ in disease-related biochemical changes, which warrant further investigation.

The major strengths of our study include a large sample size, a good proteome coverage, and evaluation of long-term stability in individuals at an age where common diseases are likely to manifest. However, some limitations must be acknowledged. One is the availability of only two measurements per individual. Longitudinal proteomics studies that include multiple time points enable a more detailed analysis of protein patterns over time and the effects of aging on circulating protein levels. With more than two time points, alternative methods, such as the intraclass correlation coefficient (ICC)^52^ could be used to determine temporal stability. For a more accurate validation of technical variability, the inclusion of duplicate samples from individuals in the AGES cohort would have been ideal. However, in the absence of such duplicates, we used reference values from the ARIC study to gain an approximate understanding of the SOMAmers‘ technical variability. It is important to note that technical variability may differ between the two cohorts due to differences in sample handling or processing protocols. Finally, as our analyses were limited to individuals with protein measurements at both visits, there is potential for selection bias. Participants who did not return for follow-up may have been in poorer health or at greater risk of disease, potentially leading to an underestimation of temporal variability or disease associations in our sample.

In conclusion, we have shown that multiple factors influence the long-term temporal stability of circulating proteins. Temporally stable and variable proteins appear to represent distinct sets of proteins with different biological functions, primarily reflecting extracellular and intracellular processes. We also demonstrated that protein temporal stability appears to reflect the mechanisms underlying proteins’ presence in circulation rather than their transcriptomic regulation. Two major contributors to temporal stability were genetic effects and disease stage, providing valuable information for biomarker discovery and precision medicine.

## Methods

### Study cohort

The AGES-Reykjavík study, conducted by the Icelandic Heart Association (IHA), is a population-based, prospective cohort study initiated in 2002 and with examinations completed in 2006^27^. The cohort comprises 5,764 Icelandic individuals aged 66 to 93, with a mean age of 76. Of these, 3,316 participants returned for a follow-up visit approximately five years later (2007–2011). Genotypic and proteomic data at two time points were available for 3,093 individuals. Information about multiple phenotypes and traits for AGES participants at both visits was available^27^, along with linked medical records. Supplementary Table 2 provides a summary of the phenotypes, abbreviations, regression models, and covariates used.

The AGES study was approved by the National Bioethics Committee in Iceland (approval number VSN-00-063), the National Institute on Aging Intramural Institutional Review Board, and the Data Protection Authority in Iceland. All participants provided informed consent before study enrollment.

### Protein data

Serum protein levels were measured at both visits using the SomaScan (*Slow Offrate Modified Aptamer Scan*, v4.1) platform, analyzing 7,569 proteins^53^. A total of 6,385 serum proteins (referred to by their gene symbols) were targeted by 7,288 SOMAmer reagents using the high-throughput SomaScan proteomic platform, after excluding 281 SOMAmers derived from mouse-human chimeras. SOMAmers are single-stranded DNA aptamers that bind to specific proteins, enabling their quantification through fluorescence intensity^53^. Samples from both visits (baseline and follow-up) were randomized, and all samples were run as a single set to minimize batch effects between visits. To ensure comparability across proteins and mitigate potential skewness in the data distribution, the Box-Cox transformation was applied, and values were standardized to a mean of 0 and a standard deviation of 1.

### Genetic data and protein quantitative trait loci (pQTLs)

AGES participants (n = 5,661) were genotyped using the Illumina Infinium Global Screening Array (GSA, n = 2,710) and Illumina hu370CNV array (n = 2,951), as previously described^6^. Genotype imputation was performed using the TOPMed reference panel^54^, yielding 11,919,579 variants after quality control and merging data from both platforms. Our team previously conducted a protein GWAS on the 5k SomaScan data (v3)^6^, and an updated analysis based on the 7k dataset (v4.1) was used in this study^35^.

Each of the 7,288 SOMAmers was tested for association with 11,919,579 variants using linear regression in PLINK 2.0, adjusting for sex, age, 10 genetic principal components, and genotyping platform. Stepwise conditional analysis was then applied with GCTA-COJO^55^ to identify independent protein quantitative trait loci (pQTLs) for each SOMAmer. The variant with the lowest p-value at each step was designated as the lead variant. Variants in strong linkage disequilibrium (LD; r² > 0.9) with the lead variant were excluded from joint analysis to avoid multicollinearity. Independent pQTLs identified by GCTA-COJO were validated by joint model testing on individual-level AGES data, from which study-wide significant signals were extracted. pQTLs with an independent lead variant within ±500 kb window of the protein-coding gene were defined as *cis*-acting; otherwise, they were classified as *trans*-acting.

### Protein temporal stability

Statistical analyses were performed in RStudio^56^ software (R version 4.3.2), unless otherwise noted. The temporal stability of each protein between baseline and follow-up visits was assessed using the Spearman correlation coefficient (SCC), hereafter referred to as the temporal SCC. Based on the distribution of these estimates, proteins were categorized into three temporal stability groups: stable, medium, and variable. Proteins with SCC higher than 0.75 were defined as temporally stable (n_prot_ = 729), whereas proteins with SCC lower than 0.40 were defined as temporally variable (n_prot_ = 681) based on suggestive guidelines for the interpretation of SCC^28^ and as they corresponded closely to the 90^th^ (0.754) and 10^th^ (0.406) percentiles of the distribution. The proteins that fell between 0.4 and 0.75 in temporal SCC were defined as having medium temporal stability.

To assess whether temporal SCC provided different insights into longitudinal variability compared to the absolute difference in protein levels between time points (Δ, delta), the delta value was calculated for each participant and protein as follows:

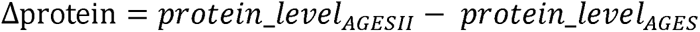

The median delta value across individuals per protein was computed and its relationship with temporal SCC estimates evaluate using the Spearman correlation coefficient. To investigate the relationship between inter- and intra-individual variability of protein levels, the inter-quartile range (IQR) value for each protein at each time point was computed to assess the inter-individual variability and compared with temporal SCC estimates.

We tested if relative protein abundance (raw protein measurements in RFU units, without transformation) differed between the temporal SCC groups. SOMAmers were measured in three dilution sets: 20% (1:5), 0.5% (1:200), and 0.005% (1:20,000), based on estimated protein concentration. The 20% dilution captured proteins in the femto- to pico-molar range, 0.5% in the nano-molar range, and 0.005% in the micro-molar range. Notably, 80% of the human proteins in the SomaScan assays were in the pico-molar range (20% dilution)^57^. To account for differences in dilution, the median raw protein value was computed for each protein from the baseline visit and adjusted using the corresponding dilution factor:

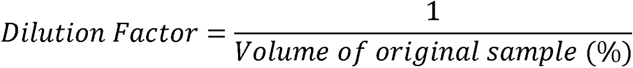

Thus, proteins in the 20% dilution set were multiplied by 1/20, while proteins in the 0.5% dilution set were multiplied by 1/0.5. To assess the relationship between temporal SCC and abundance, the distributions of median dilution-corrected RFU multiplied by the dilution factor for each protein were compared between temporal stability groups using a pairwise Wilcoxon test (two-sided). The relative protein abundance was additionally compared with external concentration values from the Human Protein Atlas^58^ (version 23.0), which provided immunoassay-based protein concentrations (pg/mL) for 569 of our 7,288 proteins, using the Spearman correlation coefficient.

For assessment of technical variability of SOMAmers, external reference values from the ARIC study were used to analyze duplicate samples from 102 individuals using the same 7k SomaScan platform (v4.1)^32^. This study estimated inter-plate stability using three different methods. For this study, the percentile variation (PV) values from the ARIC study were chosen because they demonstrated the least sensitivity to outliers and did not require the assumption of a normal distribution. Proteins with PV values more than two standard deviations above the mean were defined as having substantial technical variability. In total, 191 proteins exceeded this threshold and were classified as technically variable. These proteins were excluded from downstream pathway and functional enrichment analyses to reduce the influence of technical variability on biological interpretation of temporal stability.

### Pathway and functional enrichment analysis

Pathways and functional enrichment analyses were conducted using g:Profiler^59^ (g:GOST module) to identify potential functional differences between stable and variable proteins. A list of proteins in each temporal SCC protein group was used as an input, with the 7,097 measured proteins defined as technically stable, serving as a custom background. The significance threshold was set at the Benjamini-Hochberg false discovery rate^60^ (FDR) < 0.05. Data sources included Gene Ontology^61^ (GO), and KEGG^62^, Reactome^63^, and WikiPathways^64^. For proteins with multiple GO annotations under the same Ensembl ID, annotations with the highest number of associated GO terms were prioritized. To identify key differences between stable and variable proteins, driver GO terms were extracted and compared between the groups. Additionally, a multi-query analysis was performed, in which both lists were analyzed simultaneously using the complete set of 7,097 proteins as a background. By conducting this simultaneous analysis, the significance of enriched terms between stable and variable protein groups was directly compared, providing insights into their functional differences.

Protein class enrichment was examined using external data from the Human Protein Atlas^58^ (version 23.0), which included 5,894 proteins overlapping with the SomaScan 7k protein dataset. Fisher’s exact test^65^ was applied to identify protein classes that were significantly overrepresented in either the temporally stable or variable protein groups. Multiple testing was corrected using FDR (p_adj_ < 0.05). Additionally, using the same type of tests, enrichment for tissue-enriched genes^66^ and protein expression^67^ was assessed between temporal SCC groups.

Potential differences in protein interactions between the stable and variable protein groups were explored using both their hub status in the InWeb protein-protein interaction (PPI) network^33^ (data available for 1,443 of 3,012 proteins excluding the medium temporal stability group) and the serum protein co-regulatory network^9^ (data available for 1,940 of 3,012 proteins excluding the medium temporal stability group). This was performed using Fisher’s exact test and multiple testing correction at FDR < 0.05. Additionally, to determine whether the protein groups had differing tolerance to LoF mutations, the LoF intolerance (pLI) scores^68^ (data available for 2,924 of 3,012 proteins excluding the medium temporal stability group) were compared using two-sided Wilcoxon rank-sum test.

### Gene expression variance

Gene expression variability was investigated to determine whether it correlated with protein temporal stability. A previous study analyzed 57 publicly available RNA-sequencing datasets to examine variance in gene expression across different tissues and datasets^34^. In this study, gene expression variance was quantified as the standard deviation (SD) of gene expression levels across individuals for each gene, and each gene was ranked based on its SD values. SD-based gene expression ranks were available for 4,367 of the SOMAmers measured in AGES. These values were compared to the temporal SCC groups in AGES using a two-sided Wilcoxon rank-sum test.

### Genetic effects on temporal stability

pQTLs for the SomaScan 7k measurements in AGES (see Genetic data and protein quantitative trait loci (pQTL)) were used to assess the influence of genetic variants on the temporal SCC. A linear regression model was employed to adjust protein levels for each conditionally independent pQTL. Specifically, all independent genetic variants with genome-wide significant protein associations (p-value < 5 × 10^−8^) were included as covariates in the regression model. The residuals from this model were extracted and used as the adjusted protein values. For 716 proteins (SOMAmers) without any genetic signal, their original values were retained to ensure a balanced comparison across all proteins. Temporal SCC for the adjusted protein values was then recalculated using the same approach described above (see Protein temporal stability) and compared with the original temporal SCC estimates.

### Phenotype classification and associations with protein levels

The association between temporal stability and age-related diseases was assessed using two different models. Protein levels at baseline were tested for association with each phenotype using the full cohort, adjusted for age and sex. For prevalent phenotypes (diagnosed before or at the AGES baseline visit), a logistic regression model was used. A Cox regression model was used for incident phenotypes (diagnosed after AGES) to include the time between baseline and diagnosis, except for type 2 diabetes and chronic kidney disease, which were defined based on data from the AGESII visit and not time-to-event data (see Supplementary Table 2 for details), and thus analyzed using logistic regression. The glm() function was applied for the logistic regression analysis, and for Cox regression, the coxph() function from the survival package^69^. Multiple comparisons were controlled using FDR correction, where proteins with FDR < 0.05 were considered significantly associated with their corresponding phenotype.

### Weighted gene co-expression network analysis (WGCNA)

Weighted gene co-expression network analysis (WGCNA) was performed on protein expression levels separately for baseline and follow-up visits using the WGCNA R package^70^. To enable comparison with previously generated modules from the 5k protein dataset^9^, we set the soft-thresholding power (β) to 5 for both baseline and follow-up analyses. Co-regulatory modules were identified using dynamic tree cutting. Pathway and functional enrichment analyses were conducted for each baseline module using g:Profiler^59^, with the complete set of the 7k protein platform used as the background.

Protein modules at baseline were tested for enrichment of temporal SCC groups using two-sided Wilcoxon rank-sum test. In addition, enrichment for protein-associations with 23 diseases or phenotypes (12 prevalent and 11 incident) was tested using Fisher’s exact test, with FDR correction applied at a threshold of FDR < 0.05. Disease enrichment per module was summarized in a disease score by multiplying the number of significantly enriched diseases by the median odds ratio (OR) of those enrichments, thus accounting for both the number and strength of disease associations within each module.

Seven phenotypes (myocardial infarction [MI], coronary heart disease [CHD], congestive heart failure [CHF], atrial fibrillation [AF], Alzheimer’s disease [AD], dementia, and stroke) were further examined in relation to protein temporal stability. For each phenotype, participants were categorized into one of the four disease timeline groups:

- **Before A1**: Diagnosed before baseline
- **Between A1 and A2**: Diagnosed between baseline and follow-up
- **After A2**: Diagnosed after follow-up
- **Never**: Never diagnosed

For proteins within each baseline module, significantly associated with the respective phenotype, the temporal SCC was recalculated separately within each disease timeline group of individuals. A pairwise permutation test (10,000 permutations), shuffling timeline group labels, was used to assess whether median temporal SCC between disease timeline groups were statistically significant. The empirical p-values were corrected for multiple testing using the Benjamini-Hochberg method and FDR < 0.05 considered statistically significant.

## Acknowledgements

The authors acknowledge the contribution of the Icelandic Heart Association staff to AGES-Reykjavik as well as the involvement of all study participants. National Institute on Aging contracts N01-AG-12100 and HHSN271201200022C, the Icelandic Heart Association, and Althingi (the Icelandic Parliament) financed the AGES-Reykjavik study. IHA and Novartis have collaborated on proteomics research since 2012. This study was also funded by the Memorial fund of Helga Jonsdottir and Sigurlidi Kristjansson, the Icelandic Research Fund (2511008-051), and the Icelandic Centre for Research (2412215-1101).

## Author contributions

Conceptualization: H.K.I., Va.G. Formal Analysis: H.K.I., H.R.B., E.A.F., E.J., K.A. Resources: Vi.G., L.J.L., N.F., J.J.L., V.E. Data curation: T.A., Va.G. Writing – Original Draft Preparation: H.K.I., Va.G., Writing – Review and Editing: all authors, Supervision: Va.G., A.P., Vi.G.

