## Supplementary Materials for "Long-term temporal stability of circulating proteins in older adults"

**Affiliations**

### Supplementary Figures

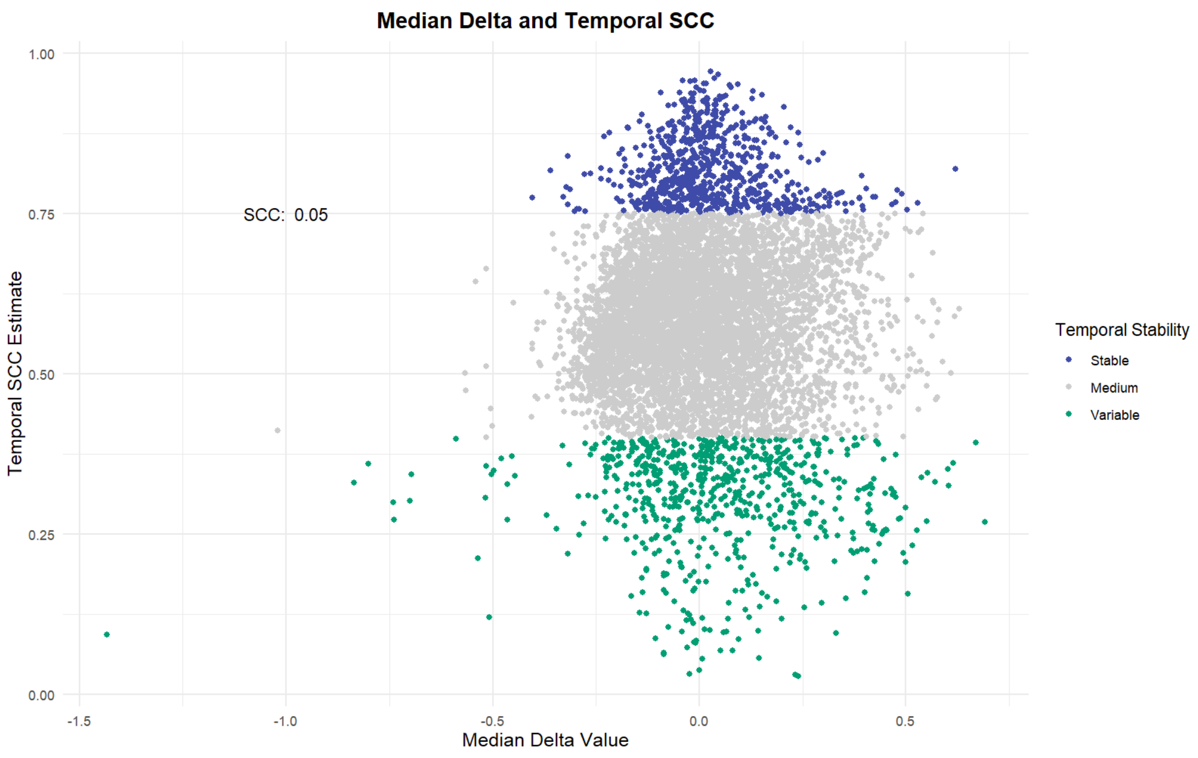

**Supplementary Figure 1**: Comparison of the median delta of each protein with its temporal SCC estimate, colored by temporal stability group. This revealed almost no correlation (SCC = 0.05, p-value = 3.90 x $10^{-6}$).

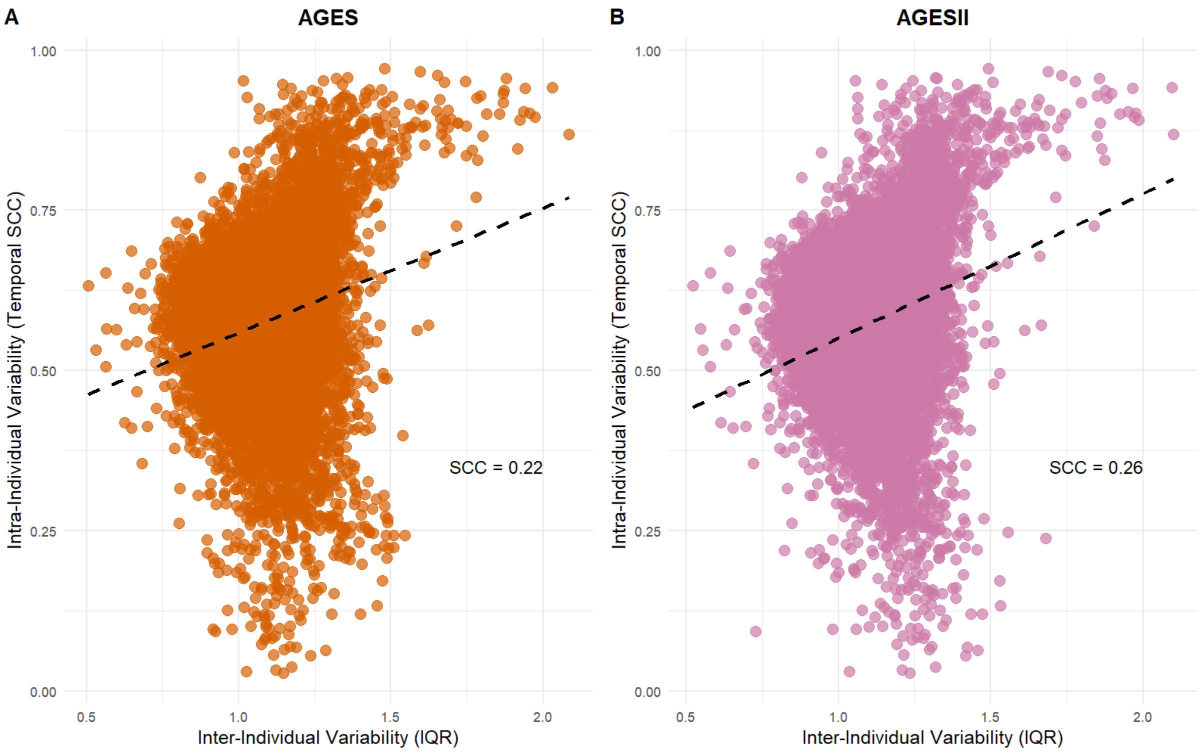

**Supplementary Figure 2: A–B)** Relationship between protein inter-individual variability (IQR) and intra-individual variability (temporal SCC) at each time point. At both visits, there was only a weak correlation between intra- and inter-individual variability (SCC at AGES = 0.22, SCC at AGESII = 0.26).

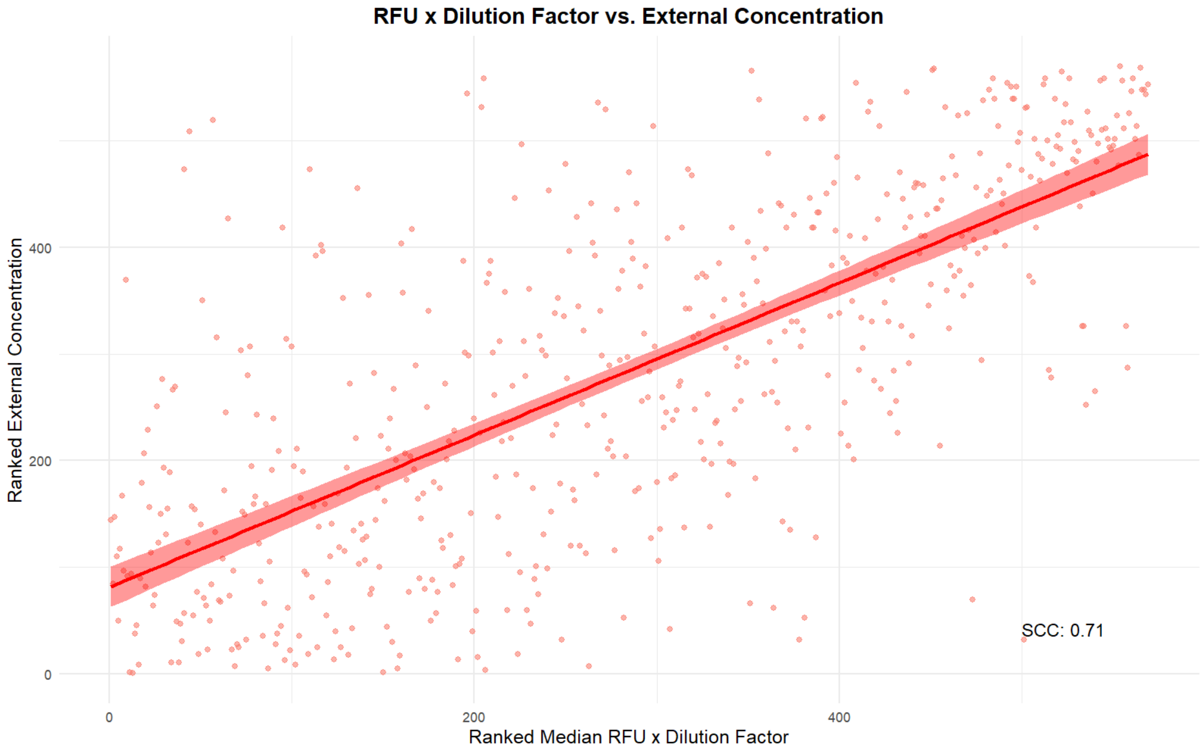

**Supplementary Figure 3:** Comparison of external concentration values for 569 proteins in blood from the Human Protein Atlas with median RFU values (multiplied by the dilution factor) from AGES showed a strong correlation (SCC = 0.71, $p < 2.2 \times 10^{-16}$). The scatterplot illustrates the ranked values and the fitted regression line with a 95% confidence interval.

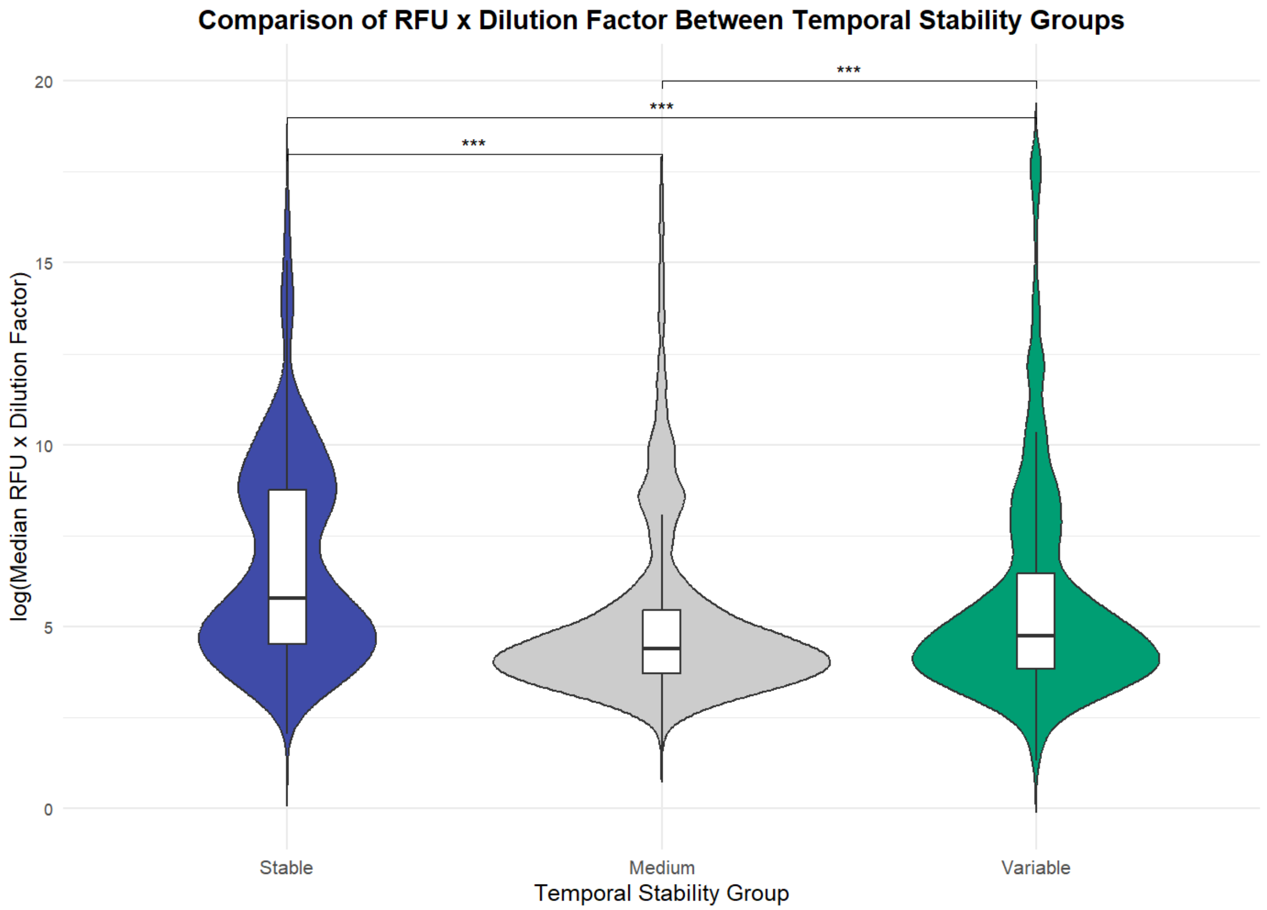

**Supplementary Figure 4:** Log-transformed median RFU values (multiplied by the dilution factor) are shown for proteins categorized as stable, medium, or variable in temporal stability. Proteins in the stable group had higher abundance compared to Medium and Variable groups (pairwise two-sided Wilcoxon tests, $p < 1 \times 10^{-3}$). Boxplots indicate median value, 25th and 75th percentiles. Whiskers extend to smallest/largest value of no further than 1.5 × interquartile range. Outliers are not shown.

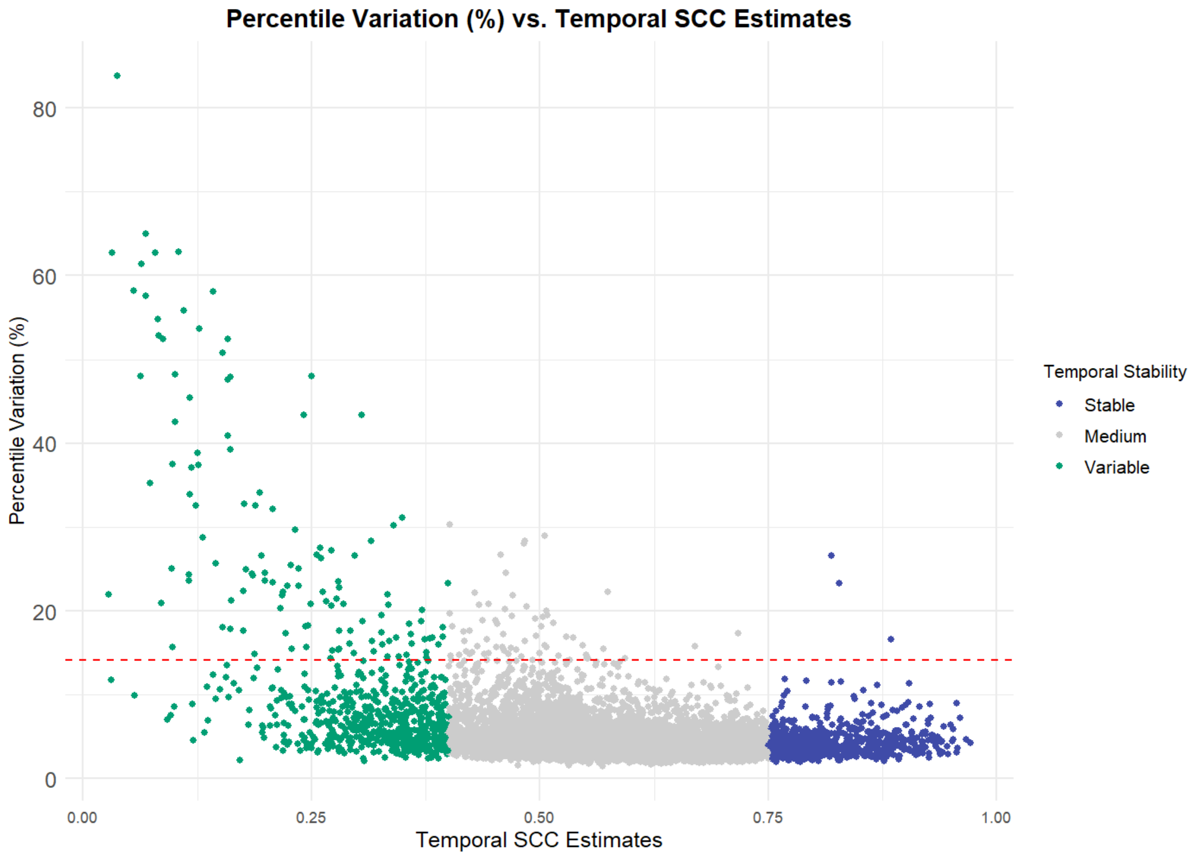

**Supplementary Figure 5:** Comparison of percentile variation (PV) as a measure of protein technical variability from ARIC with temporal SCC estimates in AGES. A cutoff of z-score > 2 was used to identify proteins with high PV, indicating high technical variability. Proteins with PV below this cutoff were considered technically stable.

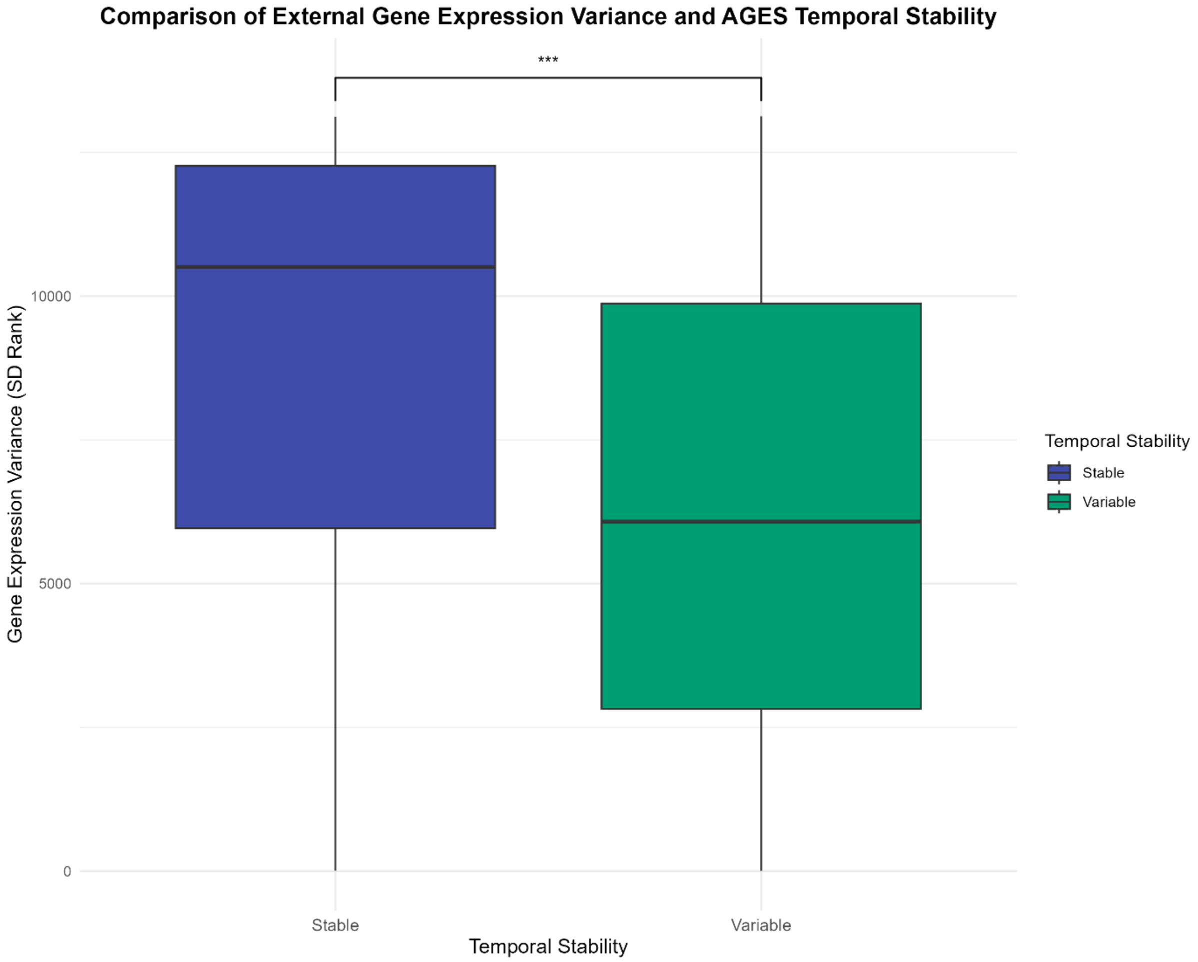

**Supplementary Figure 6:** Comparison of gene expression variance and temporal stability in proteins from AGES. Gene expression variance values (SD) from Wolf et al. (34) were compared between stable and variable protein groups. Stable proteins had significantly higher SD rank values (Wilcoxon test, two-sided, $p < 1 \times 10^{-3}$), indicating significantly more gene expression variance, while variable proteins were more likely to exhibit lower gene expression variance. Boxplots indicate median value, 25th and 75th percentiles. Whiskers extend to smallest/largest value of no further than 1.5 × interquartile range. Outliers are not shown.

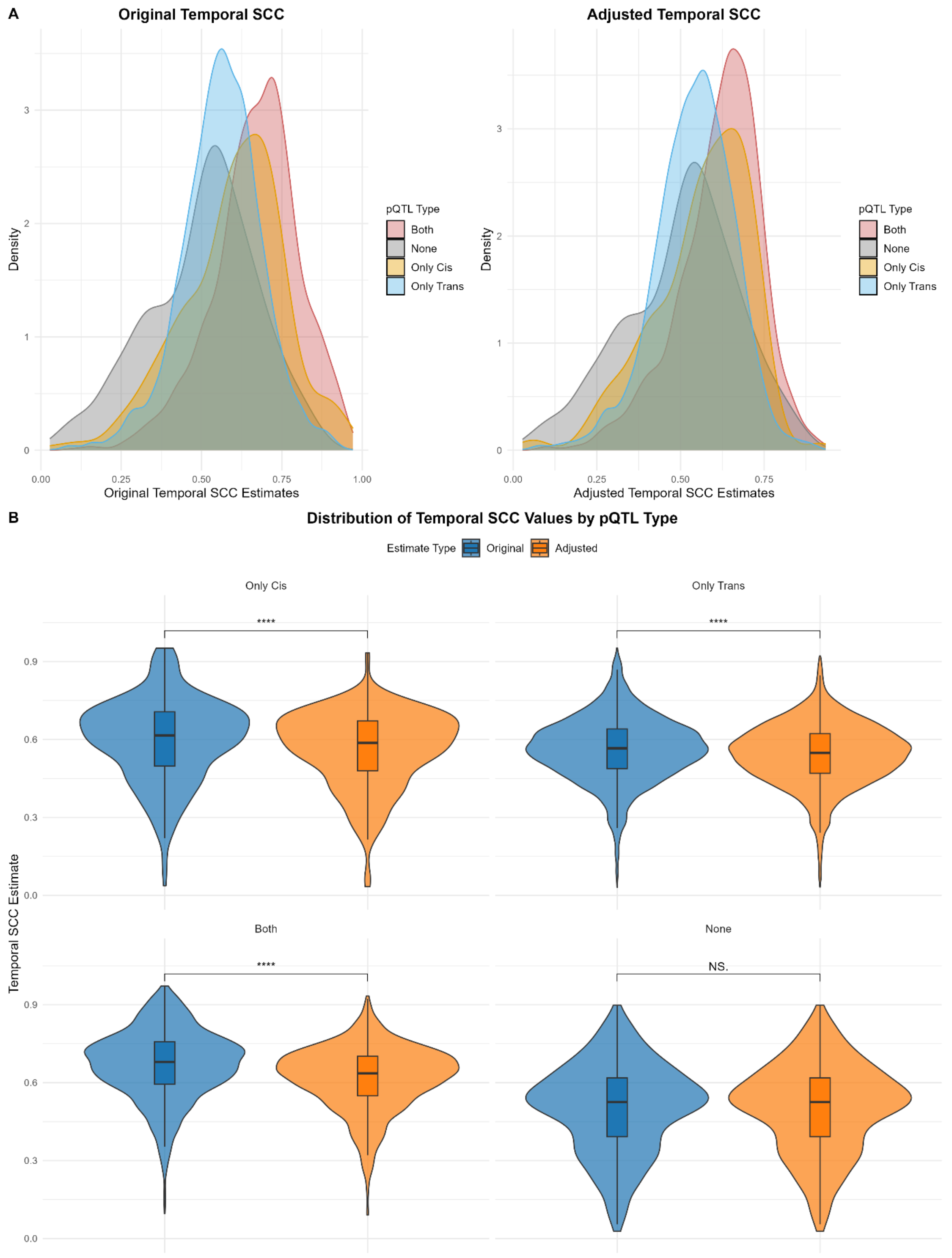

**Supplementary Figure 7: A)** Distribution of original and adjusted temporal SCC estimates by pQTL type. Proteins with higher SCC were more likely to have cis-pQTLs or both cis- and trans-pQTLs. Distributions differed significantly across pQTL groups for both original and adjusted SCC (Kolmogorov-Smirnov test, all $p < 1 \times 10^{-3}$). **B)** Change in temporal SCC after adjustment, stratified by pQTL group. Original SCC values were higher than adjusted SCC in all groups except those without any signal that remained unadjusted (paired t-test, $p < 0.0001$). Proteins with both cis- and trans-pQTLs showed the largest decreases. Boxplots indicate median value, 25th and 75th percentiles. Whiskers extend to smallest/largest value of no further than 1.5 × interquartile range. Outliers are not shown.

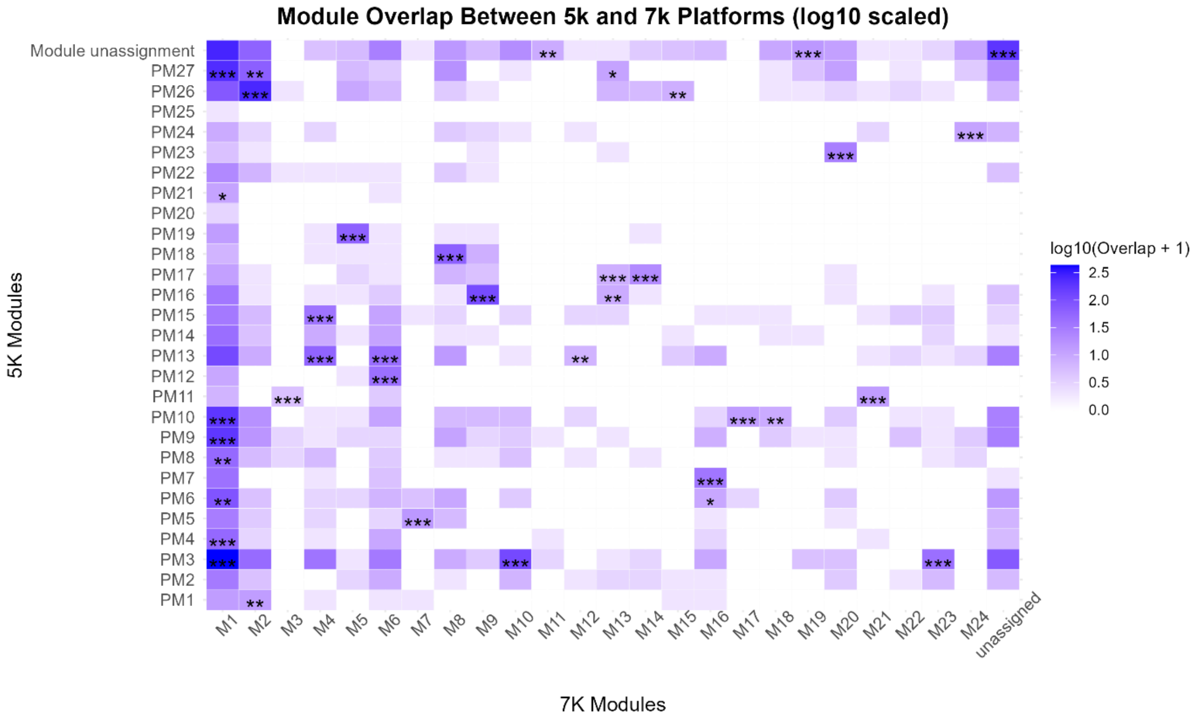

**Supplementary Figure 8:** Heatmap showing the overlap between baseline protein co-regulatory modules defined using the 5k SomaScan network (n = 5,457) and the 7k SomaScan network (n = 3,093). Tiles represent the number of shared proteins between modules, log10-transformed (log10(Overlap + 1)) for visualization. Statistical significance of overlaps (Fisher's exact test, FDR adjusted) is indicated by stars: * (FDR < 0.05), ** (FDR < 0.01), and *** (FDR < 0.001). Colors reflect log10-transformed overlap, with white indicating lower and blue higher overlap counts.
